# Identifying Novel Candidates for Re-Purposing as Potential Therapeutic Agents for Alzheimer’s disease

**DOI:** 10.1101/622308

**Authors:** C Ballard, B Creese, A Gatt, P Doherty, PT Francis, D Whitfield, A Corbett, J Corcoran, D Hanger, S Thuret, R Killick, G Williams

## Abstract

The current paper describes the identification of novel candidate compounds for repositioning as treatments for Alzheimer’s disease (AD) from the CMAP library. Candidate compounds were identified based on inverse correlation with transcriptome signatures developed from meta-analyses of Alzheimer RNA expression studies using the SPIED platform. The 78 compounds with a significant inverse correlation were taken forward into an in vitro programme using 6 well validated screening assays relevant to potential treatment targets in AD. Nineteen pf the compounds were hits in at least 2 of these assays. A description of each of these compounds is presented.

## Background

Alzheimer’s disease (AD) is a devastating progressive neurodegenerative disease affecting more than 45 million people worldwide (Livingston et al 2017). Two classes of symptomatic pharmacological treatments, memantine and cholinesterase inhibitors, confer modest but important short term benefits (Birks and Harvey 2018, Howard et al 2012)) but treatments with a more sustained impact are urgently needed. Despite considerable efforts, there are still no effective disease modifying treatments for AD, and there have been no new licensed therapeutics for 20 years. A number of candidate disease-modifying compounds for AD, mainly focussed on single specific disease targets, have failed in clinical trials over the last five years, with some recent high profile failures (Egan et al 2019, Honig al 2018)). Robust innovative approaches are urgently needed to re-invigorate drug discovery for AD and identify compounds with multiple impacts on key pathways that will modify disease course and make a real differences to the lives of millions of people worldwide.

Drug repositioning offers a novel, attractive and cost-effective alternative (Corbett et al 2012). It has been the basis of successful therapies in many clinical areas including diverse areas such as cancer (Heckman-Stoddard et al 2017) smoking cessation (Evins et al 2019) and Parkinson’s disease (Hubsher et al 2012). The established safety of candidate compounds is a huge advantage, and the time and cost associated with bringing these compounds to clinical trials is massively reduced (Corbett et al 2012).

The overarching aim this work is to identify novel potential candidates for re-purposing as treatments for AD based on mRNA expression profiles.

## Method

### Overview

The work was undertaken in 3 key stages: (1) Developing a transcript signature for early and mild Alzheimer’s disease based on published mRNA expression studies, (2) using this signature to identify the most promising novel compounds for repurposing as potential new treatments for Alzheimer’s disease based on the signatures compounds within the connectivity map (CMAP library) and related databases (3), to evaluate the “hits” in a broad range of Alzheimer related in-vitro assays examining amyloid and non-amyloid related potential treatment targets.

### AD associated transcriptional profiles

Transcriptional profiles associated with AD were obtained from the NCBI GEO database [Barrett, T. et al. Nucleic Acids Res. 2007, 35, D760-5.] by performing queries for series containing samples derived from post-mortem AD patient brains for various stages of the disease, together with mouse models of the disease. Profiles were defined as 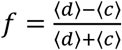, where the brackets indicate averages of the control (c) and disease (d) samples. The statistical significance is measured by a student’s t-test and those folds falling below the 95% confidence interval were dropped. The human disease versus control AD set comprises 21 profiles derived from 14 series (NCBI GEO accession: GSE84422, GSE37263, GSE36980, GSE39420, GSE36980, GSE1297, GSE29378, GSE48350, GSE15222, GSE26972, GSE37264, GSE28146, GSE5281, GSE13214). Cognitive decline was based on decline in MMSE represented by two profiles from two independent series and CDR profiles from one series. Murine model profiles were gathered from 5xFAD and 3xTG datasets giving seven profiles from three series (NCBI GEO accession: GSE50521, GSE119756, GSE101144, GSE77574) for the 5xFAD set and nine profiles from eight series (NCBI GEO accession: GSE31624, GSE15128, GSE36237, GSE92926, GSE60460, GSE60911, GSE36981, GSE35210) for the 3xTG set. BRAAK stage progression profiles (NCBI GEO accession: GSE1297, GSE84422, GSE48350, GSE106241) were generated with a linear mixed model analysis, by fitting the gene expression level across the samples in the series to a linear function of the BRAAK stage with categorical calls on cell type and gender as covariates. The resulting residual correlation Z score for gene expression against BRAAK stage constituted the BRAAK profile. 12 profiles were extracted from four series, with six following full BRAAK progression from level 0 to level 6 and six profiles corresponding to mild BRAAK pathology, level 0 to level 3. Similar profiles were generated for psychiatric measures MMSE and CDR (NCBI GEO accession: GSE48350, GSE1297, GSE84422). In the case of the MMSE profile the regression signs were reversed as MMSE scores decrease with disease progression.

### Compound candidate list

Each of the AD associated profiles was queried against CMAP and LINCS drug transcriptional profile databases. Each compound is assigned a rank based on the top anti-correlation across the profiles in each category: AD, BRAAK, BRAAK/mild AD, COG, 5FAD, 3TG. Table 1 lists the candidate compounds together with their top ranks in the AD profile sets. There is a good agreement between CMAP and LINCS.

**Table 1:** Compounds which were “Hits” on at Least 2 *in vitro* Assays.

| Compound | Source | CMAP<br>Correlation | A-beta cell death | Dendritic spine<br>loss | P-tau<br>reduction<br>(CHO<br>model) | APP-mediated<br>Wnt<br>restoration | DKK<br>Blocking | Neurogenesis |
| --- | --- | --- | --- | --- | --- | --- | --- | --- |
| Imatinib | LINCS | -0.18<br>(Early AD) | p=0.22<br>ES 27.5% | p=0.0267 |  | p=0.02 |  |  |
| Mitoxantrone | CMAP | -0.27<br>(Mouse sig) |  | p=0.012 | p<0.05, d>1 | p=0.008 |  |  |
| Kenpaullone | LINCS | -0.2<br>(Mod AD) |  |  | p<0.05, d>1 | p=0.00 | p=0.00 |  |
| Tacrine | CMAP | -0.21<br>(Early AD) | p=0.007<br>ES 35.9% |  | p<0.05, d>1 |  |  |  |
| Apigenin | CMAP | -0.09<br>(Mouse sig) | p=0.0001<br>ES 72.5% |  | p<0.05, d>1 |  |  |  |
| Chlortetracyclin | CMAP | -0.11<br>(Mouse sig) | p=0.003<br>ES 32.1% |  |  | p=0.00 |  |  |
| Thiostrepton | CMAP | -0.16<br>(Early AD) | p<0.000<br>ES 46.8% |  |  | p=0.05 |  | Flagged as possible<br>negative impact |
| Trioxsalen | NMAP | -0.12<br>(Early AD) | p=0.0003<br>ES 41% |  |  | p=0.00 |  |  |
| Famotidine | NMAP | -0.17<br>(Early AD) | p=0.14<br>ES 23.8% |  |  |  | p=0.001 |  |
| Cyanocobalamin | CMAP | -0.29<br>(Early AD) | p=0.003<br>ES 39.1% |  |  |  |  |  |
| Doxazosin Mesylate | NMAP | -0.16<br>(Early AD) | p=0.12<br>ES 51% |  |  |  |  |  |
| Luteolin | CMAP | -0.09<br>(Mouse sig) |  | p=0.0079 | p<0.05, d>1 |  |  |  |
| Hexestrol | NMAP | -0.12<br>(Mouse sig) |  | p=0.0086 | p<0.05, d>1 |  |  |  |
| Metformin | CMAP | -0.8<br>(PD) |  |  | p<0.05, d>1 |  |  |  |
| Yohimbine | CMAP | -0.11<br>(PD) |  |  | p<0.05, d>1 |  |  |  |
| Pyrantel | CMAP | -0.17<br>(Early AD) |  |  | p<0.05, d>1 |  |  |  |
| Cotinine | CMAP | -0.58<br>(PD) |  |  |  | p=0.00 | p=0.034 |  |
| Genistin | LINCS | -0.17<br>(Mod AD) |  |  |  | p=0.00 | p=0.00 |  |
| Nabumetone | CMAP | -0.30<br>(PD) |  |  |  | P=0.03 | P=0.000 |  |

**Table 2.**
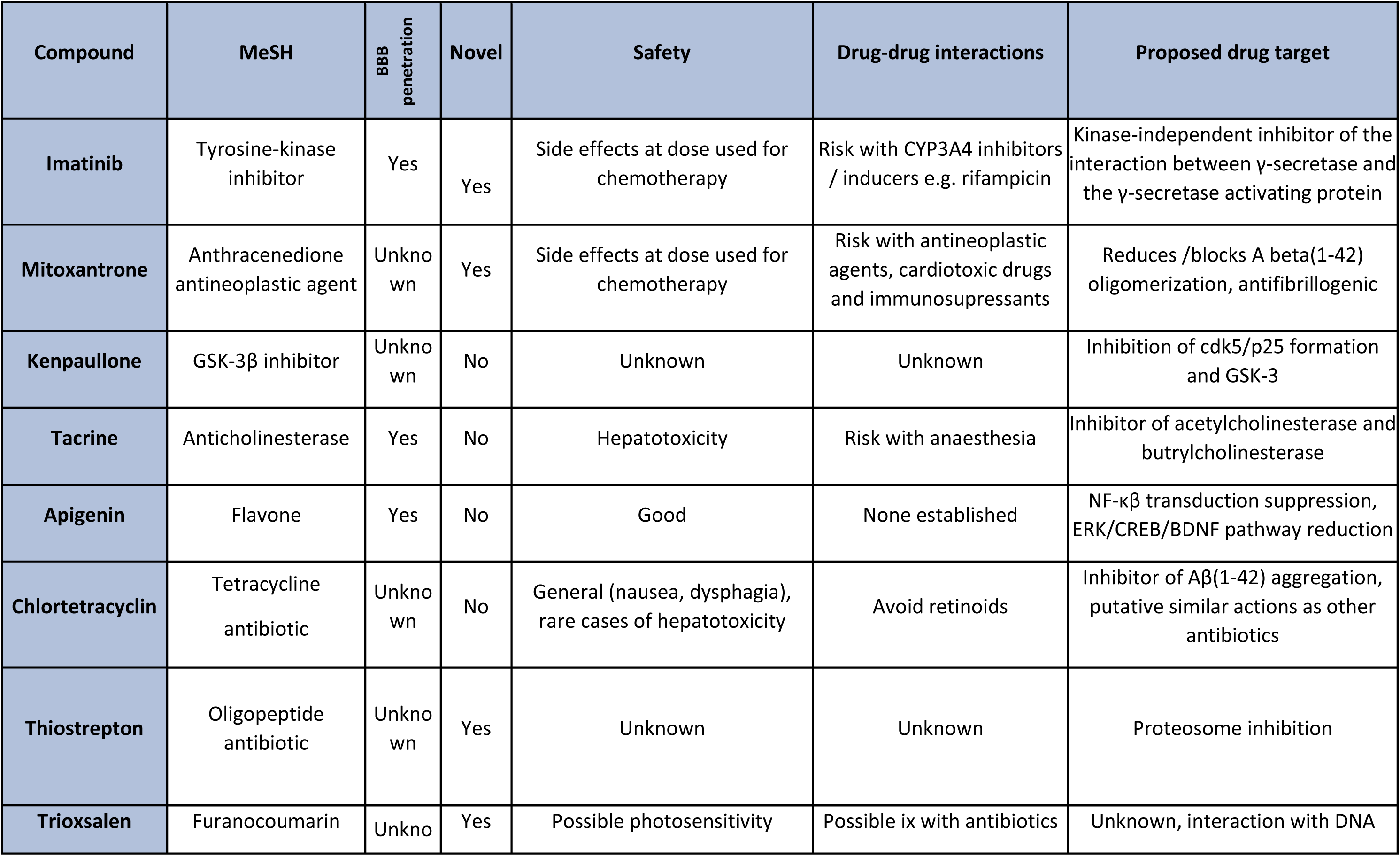
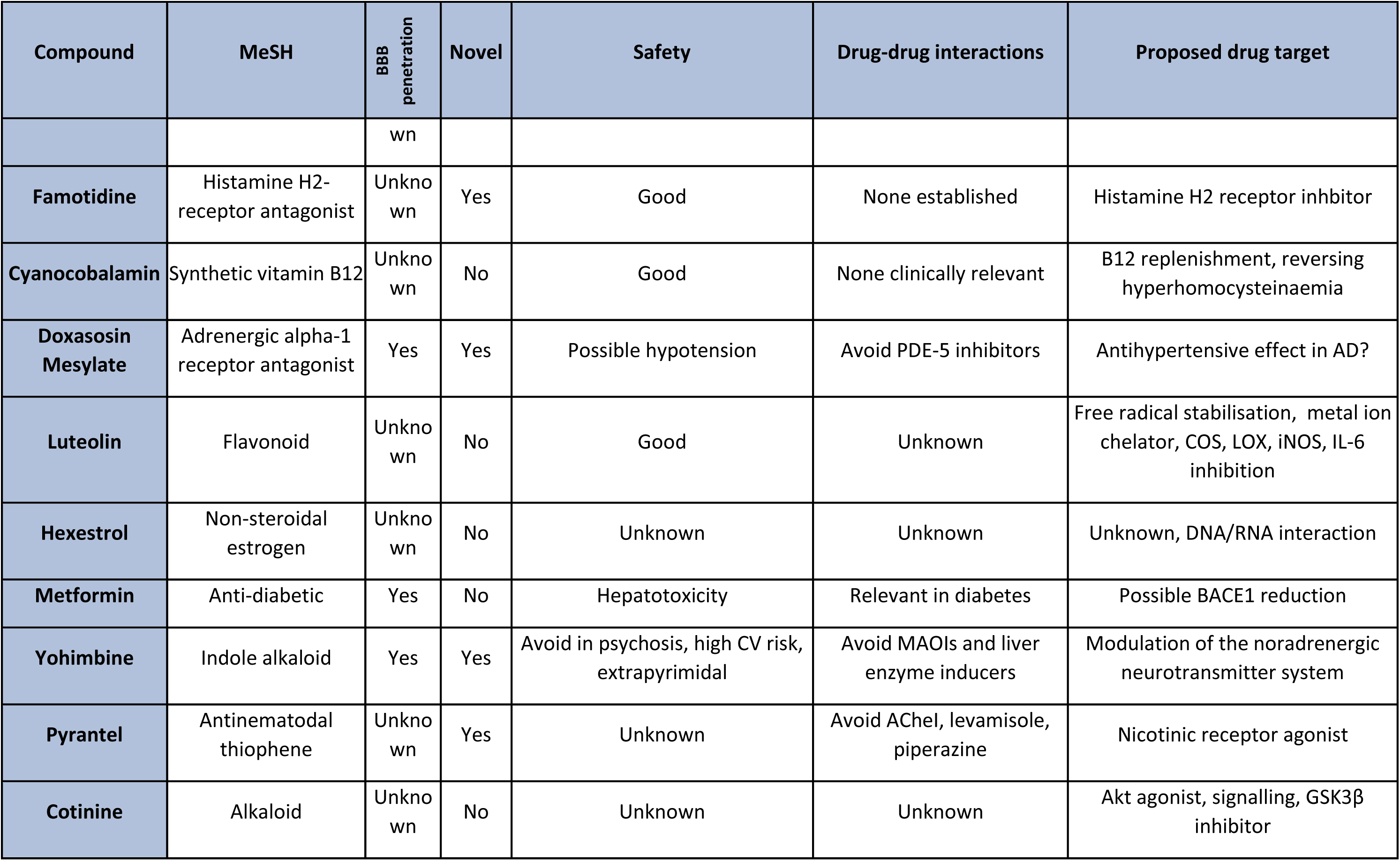
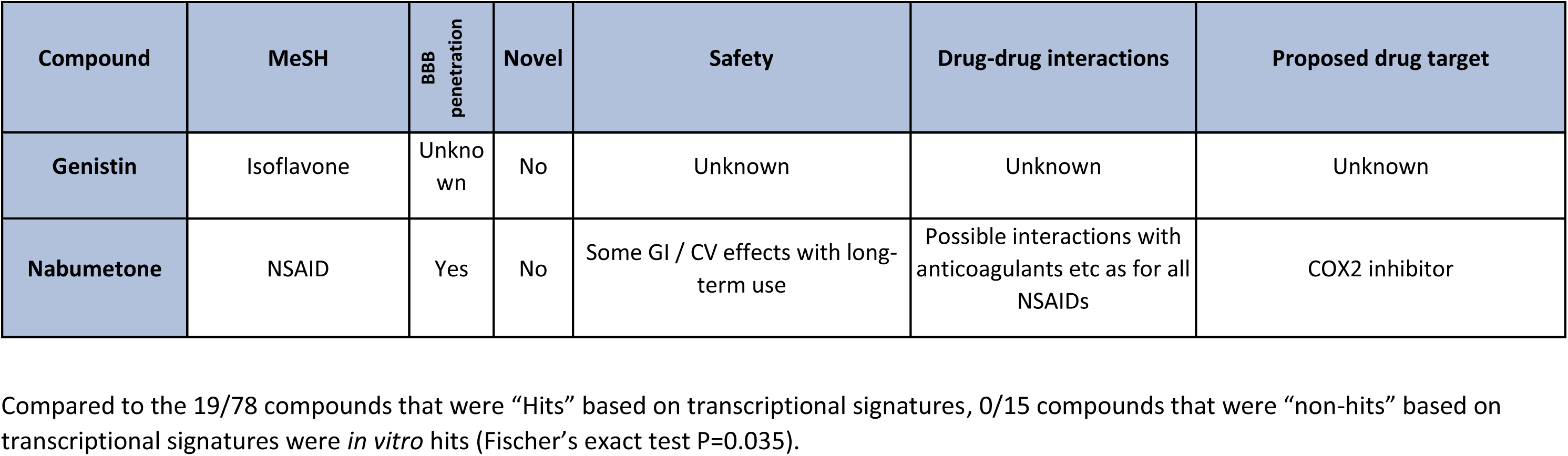
Characteristics of compounds which were hit an at least 2 in vitro assays.

### CMAP and LINCS databases

CMAP data was downloaded from the Broad connectivity map site (www.broadinstitute.org/connectivity-map-cmap). This consisted of probe sets for each sample ranked according to expression level relative to batch control. The data consists of 6,100 samples covering 1,260 drugs and four cell types. The relative probe expression ranks, defined as 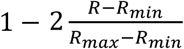, where *R* in the rank of a given gene’s expression change (*R*_*max*_ being the highest and *R*_*min*_ being the lowest ranks), were averaged over replicates ingnoring cell type and filtered based on significance using a one sample student’s t-test. For genes with multiple probes the probe with the largest significant change was mapped to the gene. This resulted in a unique profile for each drug in CMAP. The compound data can be queried through SPIED (Williams 2013).

LINCS data was based on the GEO depositories GSE92742 and GSE70138, which provide Z-scores for compound versus control treatment data. Replicates were combined and statistically filtered giving a dataset unique for compound/cell type pairings.

### Evaluation of the neuroprotective efficacy of the best candidate drugs in vitro

#### The selected assays included

Mouse cortical neuron survival in presence of a-beta: This is an established assay for analysis of neuronal cell death following insult by a-beta and followed well established protocols (Goncalves et al 2013). In brief, mouse cortical neurons were treated with oligomeric amyloid and candidate compound in 96-well plate format for three days. The outcome measure was cell survival using Tunel measurement. Compounds conferring a significant improvement in cell survival (p<0.05) were considered to be “hits”.

Dendritic Spine Loss: This assay evaluated the ability of compounds to block Aβ-driven spine loss will maintain neuronal viability and potentially protect cognition in vivo. Primary rat neurons cultured for 24 days and transfected with EGFP.30 minutes pre-treatment with compound, followed by four-hour incubation with 20uM Aβ_25-35_ oligomers. Fixation was undertaken for immunocytochemistry. Antibodies against PSD-95 and synapsin were imaged through confocal microscopy. Dendritic spine morphology scores were rated for: (1) Identified valid cell bodies, (2) Average length neurites / neuron; (3) No. branch points / neuron; (4) No. PSD-95 puncta / neurite; (5) No. Synapsin-1 puncta / neurite; (6) No. synapses / neuron calculated as overlap of PSD-95 and synapsin-1. Compounds conferring significant benefits in the combined spine loss score (p<0.05) were considered to be “hits” (Killick et al 2018)

APP-mediated wnt restoration: The wnt pathway is known to be disrupted in AD pathology, providing a potential target for AD treatments. Luciferase-based Wnt reporter gene assay was performed in Human HEK293T neuron-like cells in a 96-well format. TOPflash and FOPflash genes were used as reporters. Cells were treated overnight with the reporter, wnt3a and Dkk1. Each replicate was n = 8. Benefit was evaluatd based on restoration of canonical wnt signalling activity as measured by luciferase output. Compounds conferring significant benefit with respect to wnt restoration (p<0.05 and Effect Size of 20%) were considered to be “hits” (Killick et al 2018)

<u>DKK:</u> Blocking Aβ induction of Dkk1 blocks many important neurotoxic effects of Aβ, and therefore forms an important potential treatment target for AD. Human HEK293T cells were treated overnight following the same protocol as the APP-mediated wnt restoration assay described above. Each replicate was n = 8. The evaluated outcome was Dkk1 protein expression determined by an ELISA-based measure of Dkk1 levels. Compounds conferring significant dkk blocking activity (p<0.05 and Effect Size of 20%) were considered to be “hits” (Killick et al 2018).

Reduction of total and phosphorylated tau: Production and phosphorylation of tau is an established measurement of AD pathology and a key potential treatment target. CHO cells stably expressing Tau35 were treated for four hours with each compound in a 96-well format. Cells were fixed and stained for total tau, phosphorylated tau (Ser-396) and actin. Each replicate was n = 8. Total and phosphorylated tau as described previously (ref). Compounds achieving a significant reduction in tau phosphorylation (p<0.05 and Effect Size of 20% reduction) were considered to be “hits” (Noble et al 2009).

Neurogenesis (cell proliferation and differentiation): This assay provides a model for studying molecular mechanisms regulating neurogenesis, a major question in potential AD treatment candidates. HPC03A/07 human hippocampal stem cell lines were treated for for 48 hours (proliferation assay) and seven days (differentiation assay) respectively in a 96-well format. Cells were stained for stem-cell markers (Sox2, Nestin), proliferation (Ki67), apoptotic cell death (CC3), immature (Dcx) and mature (Map2) neurons and astrocytes (S100beta). The assay was analysed based on the percentage change from baseline with treatment on six readouts: (1) cell number, (2) cell proliferation, (3) Neurogenesis [(3a) neuroblasts, (3b) neurons, (3b) neurite outgrowth and (4) cell death. Compounds which increased proliferation of hippocampal progenitor cells or cells expressing stemness markers (Sox2 / Nestin), increased differentiation of hippocampal progenitor cells and decreased apoptotic cell death p<0.05 on each of these measurements) were considered to be “hits” (Thuret et al 2013).

For all assays, the internal controls were amyloid alone, vehicle alone and compound plus vehicle.

## Results

### CMAP, LINCS and NMAP

Seventy-eight compounds from the 2 databases achieved an inverse correlation with a p value <0.05 with at least one of the 3 AD signatures.

**Figure 1.**
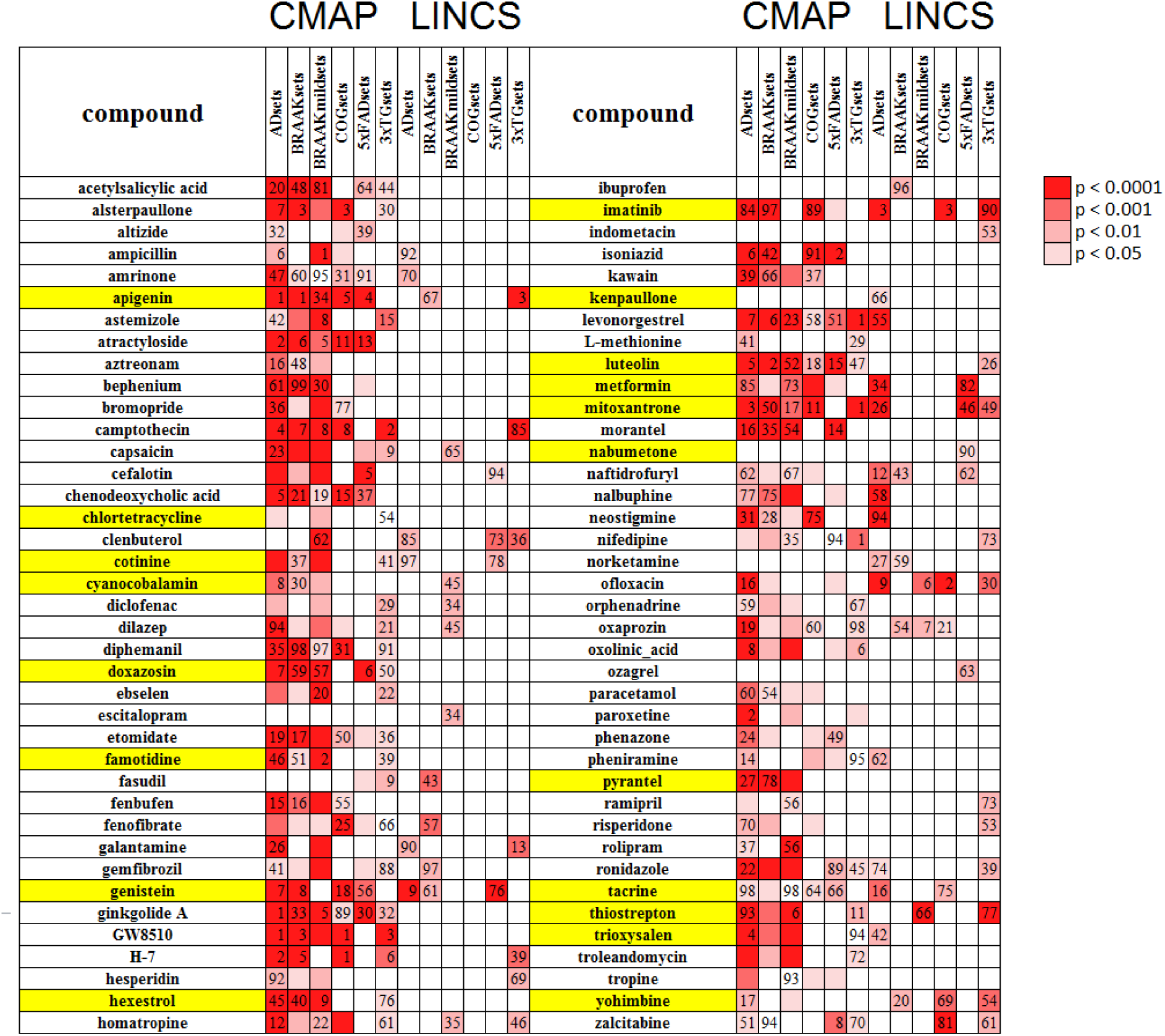
Compounds with P<0.05 inverse correlation between AD Signatures and Drug Signatures from CMAP and/or LINCS. Figure 1 footnote. The anti-correlating ranks of AD associated profiles against the CMAP and LINCS databases. AD profiles were based on: post mortem overt diseases versus control samples (ADsets); expression changes correlating with full BRAAK progression; expression changes correlating with mild BRAAK (<=3) progression; cognitive decline, based on Mini-Mental State Examination (MMSE) and *Clinical Dementia Rating (*CDR); sets of 5xFAD mouse brain samples; sets of 3xTG mouse model brain samples. The numbers given are the ranks of the given compound for the given query, where the rank is for the anti-correlation score. The drugs highlighted in yellow are those showing positive activity in at least two in vitro neuroprotection assays.

### In Vitro Hits

Of the 78 compounds examined in the in vitro assays, 19 compounds fulfilled the criteria of reaching significance on two independent assays. The data for these compounds and a summary of their results in the in vitro studies are presented in table 1.

## Discussion

The 3 stage approach in the current study developed a series of mRNA expression signatures for AD, used these signatures to identify compounds which were inversely associated with this signature in the CMAP and LINCS databases and then evaluated these compounds in a series of well validated in vitro assays focussing on target treatment mechanisms relevant to AD. Seventy eight compounds were identified which produced signatures which were significantly inversely correlated with the signatures for mild AD, moderate-severe AD or the signature of the 5Xfad AD mouse. Of these compounds, 19 (24%) were “hits” in at least two of the in vitro assays.

Firstly, the proportion of “CMAP hits” that were “hits” in the in vitro studies was significantly higher than the proportion of hits from CMAP non-hits (24$% vs 0%, FET P=0,035). In addition, the 24% proportion of compounds identified as hits in in vitro studies compares favourably to reported drug discovery work using high throughput screening (refs). This suggests that mRNA expression signatures are potentially a very useful tool to triage compound libraries and increase the likelihood of identifying compounds with positive disease related effects in in vitro studies.

Eight of the “hits” are novel candidates, 3 which are known to be brain penetrant and 5 with properties indicating a likelihood of brain penetrance. Other hits confirm candidate drugs already highlighted as potential candidates for re-purposing, such as metformin and others highlight novel drugs within classes already highlighted as potential candidate treatments, such as the cox-2 inhibitor nabumetone and several flavonoids. Overall a number of potentially promising candidates for re-purposing have been identified and merit further evaluation.

## References

Anacker C, Cattaneo A, Luoni A, Musaelyan K, Zunszain PA, Milanesi E, Rybka J, Berry A, Cirulli F, Thuret S, Price J, Riva MA, Gennarelli M, Pariante CM. Glucocorticoid-related molecular signaling pathways regulating hippocampal neurogenesis. Neuropsychopharmacology. 2013 Apr;38(5):872–83. doi:10.1038/npp.2012.253. Epub 2012 Dec 6. PMID:23303060

Barrett T, Troup DB, Wilhite SE, Ledoux P, Rudnev D, Evangelista C, Kim IF, Soboleva A, Tomashevsky M, Edgar R. NCBI GEO: mining tens of millions of expression profiles--database and tools update. Nucleic Acids Res. 2007 Jan;35(Database issue):D760-5.PMID17099226

Birks JS, Harvey RJ. Donepezil for dementia due to Alzheimer’s disease. Cochrane review 2018

Corbett A, Pickett J, Burns A, Corcoran J, Dunnett SB, Edison P, Hagan JJ, Holmes C, Jones E, Katona C, Kearns I, Kehoe P, Mudher A, Passmore A, Shepherd N, Walsh F, Ballard C. Drug repositioning for Alzheimer’s disease. Nat Rev Drug Discov. 2012 Nov;11(11):833–46. doi:10.1038/nrd3869.

Egan MF, Kost J, Voss T, Mukai Y, Aisen PS, Cummings JL, Tariot PN, Vellas B, van Dyck CH, Boada M, Zhang Y, Li W, Furtek C, Mahoney E, Harper Mozley L, Mo Y, Sur C, Michelson D. Randomized Trial of Verubecestat for Prodromal Alzheimer’s Disease. N Engl J Med. 2019 Apr 11;380(15):1408–1420. doi:10.1056/NEJMoa1812840

Elliott C, Rojo AI, Ribe E, Broadstock M, Xia W, Morin P, Semenov M, Baillie G, Cuadrado A, Al-Shawi R, Ballard CG, Simons P, Killick R. A role for APP in Wnt signalling links synapse loss with β-amyloid production. Transl Psychiatry. 2018 Sep 20;8(1):179. doi:10.1038/s41398-018-0231-6.

Evins AE, Benowitz NL, West R, Russ C, McRae T, Lawrence D, Krishen A, St Aubin L, Maravic MC, Anthenelli RM. Neuropsychiatric Safety and Efficacy of Varenicline, Bupropion, and Nicotine Patch in Smokers With Psychotic, Anxiety, and Mood Disorders in the EAGLES Trial. J Clin Psychopharmacol. 2019 Mar/Apr;39(2):108–116. doi:10.1097/JCP.0000000000001015.

Goncalves MB, Clarke E, Hobbs C, Malmqvist T, Deacon R, Jack J, Corcoran JP. Amyloid inhibits retinoic acid synthesis exacerbating Alzheimer disease pathology which can be attenuated by an retinoic acid receptor agonist. Eur J Neurosci. 2013 Apr;37(7):1182–92. doi:10.1111/ejn.12142. Epub 2013 Feb 4.

Heckman-Stoddard BM, DeCensi A, Sahasrabuddhe VV, Ford LG. Repurposing metformin for the prevention of cancer and cancer recurrence. Diabetologia. 2017 Sep;60(9):1639–1647. doi:10.1007/s00125-017-4372-6.

Honig LS, Vellas B, Woodward M, Boada M, Bullock R, Borrie M, Hager K, Andreasen N, Scarpini E, Liu-Seifert H, Case M, Dean RA, Hake A, Sundell K, Poole Hoffmann V, Carlson C, Khanna R, Mintun M, DeMattos R, Selzler KJ, Siemers E. Trial of Solanezumab for Mild Dementia Due to Alzheimer’s Disease. N Engl J Med. 2018 Jan 25;378(4):321–330. doi:10.1056/NEJMoa1705971.

Howard R, McShane R, Lindesay J, Ritchie C, Baldwin A, Barber R, Burns A, Dening T, Findlay D, Holmes C, Hughes A, Jacoby R, Jones R, Jones R, McKeith I, Macharouthu A, O’Brien J, Passmore P, Sheehan B, Juszczak E, Katona C, Hills R, Knapp M, Ballard C, Brown R, Banerjee S, Onions C, Griffin M, Adams J, Gray R, Johnson T, Bentham P, Phillips P. Donepezil and memantine for moderate-to-severe Alzheimer’s disease. N Engl J Med. 2012 Mar 8;366(10):893–903. doi:10.1056/NEJMoa1106668.

Hubsher, G., Haider, M. & Okun, M. S. Amantadine: the journey from fighting flu to treating Parkinson disease. Neurology 78, 1096–1099 (2012).

Livingston G, Sommerlad A, Orgeta V, et al. Dementia prevention, intervention, and care. The Lancet 2017; 390(10113):2673–734.

Noble W, Garwood C, Stephenson J, Kinsey AM, Hanger DP, Anderton BH. Minocycline reduces the development of abnormal tau species in models of Alzheimer’s disease. FASEB J. 2009 Mar;23(3):739–50. doi:10.1096/fj.08-113795. Epub 2008 Nov 11.

Seybi KI, Schuman R, Ni J, Huang MM, MichaeIis ML, Glicksman MA. Identification of Small Molecule Inhibitors of Beta-Amyloid Cytotoxicity through a Cell-Based High-Throughput Screening Platform Journal of Biomolecular Screening 2008:870–878

Williams G. SPIEDw : a searchable platform-independent expression database web tool. BMC Genomics. 2013 Nov 7;14:765. doi:10.1186/1471-2164-14-765. PMID:24199845

